# Polyvalent mRNA vaccination elicited potent immune response to monkeypox surface antigens

**DOI:** 10.1101/2022.11.29.518427

**Authors:** Zhenhao Fang, Paul A. Renauer, Kazushi Suzuki, Sidi Chen

**Affiliations:** Department of Genetics, Yale University School of Medicine, New Haven, CT, USA; System Biology Institute, Yale University, West Haven, CT, USA; Center for Cancer Systems Biology, Yale University, West Haven, CT, USA; Immunobiology Program, Yale University, New Haven, CT, USA; Molecular Cell Biology, Genetics, and Development Program, Yale University, New Haven, CT, USA; MD-PhD Program, Yale University, New Haven, CT, USA; Comprehensive Cancer Center, Yale University School of Medicine, New Haven, CT, USA; Stem Cell Center, Yale University School of Medicine, New Haven, CT, USA; Center for Biomedical Data Science, Yale University School of Medicine, New Haven, CT, USA; Wu-Tsai Institute, Yale University, New Haven, CT, USA; Center for RNA Science and Medicine, Yale University, New Haven, CT, USA

**Keywords:** Monkeypox, mRNA vaccine, lipid nanoparticle, MPXVac-097, vaccinia virus, modified vaccinia Ankara, A35R, E8L

## Abstract

The soaring global monkeypox cases lead to a surge in demand for monkeypox vaccine, which far exceeds the supply. mRNA vaccine has achieved great success in prevention of coronavirus disease and holds promise against diverse pathogens. In this study, we generate a polyvalent lipid nanoparticle (LNP) mRNA vaccine candidate for monkeypox virus (MPXV) and evaluate its immunogenicity in animal models. This polyvalent MPXV mRNA vaccine candidate, MPXVac-097, encodes five 2022 MPXV targets that are important surface antigens. Three-dose (prime-boost-booster) MPXVac-097 vaccination elicits strong antibody response to A35R and E8L antigens, moderate response to M1R, but not B6R or A29, highlighting the differences in immunogenicity. Bulk T cell receptor (TCR) sequencing reveals preferential usage of VJ combinations and clonal expansion of peripheral T cells after MPXVac-097 vaccination. These data demonstrate initial feasibility of developing MPXV mRNA vaccine and pave the way for its future optimization.

## Main

To date (August 28, 2022), the ongoing monkeypox outbreak has led to more than 47,000 confirmed cases and 12 deaths in over 90 countries. Because of its fast spread, world health organization declared the monkeypox outbreak a global public-health emergency on July 23^rd^. A retrospective study recently reported evidence of asymptomatic or undiagnosed monkeypox cases during early stage of monkeypox outbreak suggesting that asymptomatic cases fueled this epidemic, and symptom-based testing and quarantining may not suffice to contain monkeypox outbreak^1^. In addition to testing and quarantining, an efficient measure to contain monkeypox virus (MPXV) transmission is the use of monkeypox vaccines for high-risk groups and close contacts of patients^2^. As the monkeypox outbreak grows rapidly, the demand for monkeypox vaccines surges drastically and many countries are facing the vaccine supply shortage.

Vaccination is the ultimate approach to prevent infectious disease outbreak from developing into a global pandemic. Replication-deficient modified vaccinia ankara (MVA) vaccine, JYNNEOS (commercial name in U.S., IMVANEX in Europe and IMVAMUNE in Canada) is the approved and preferred vaccine for monkeypox due to less side effects and contraindications compared to ACAM2000, which is a replication-competent vaccinia vaccine also available for use against monkeypox^2-4^. The monkeypox vaccine in short supply is JYNNEOS, while ACAM2000 supply is abundant. However, the ACAM2000 vaccination is associated with unexpectedly high rate of myocarditis and pericarditis^5^, and can cause severe illness in immunocompromised patients. The authorization of JYNNEOS and ACAM2000 is based on 1) a phase 3 trial assessing neutralizing antibody response to vaccinia virus in vaccinees of MVA and ACAM2000^6^; and 2) a MPXV challenge experiment in macaques showing 100% protection after two doses of MVA or one dose of ACAM2000 injection^7^. Although no data is currently available on the efficacy of either vaccine against 2022 MPXV in human, a series of clinical trials and cohort studies are underway to evaluate MVA vaccine’s efficacy, safety and protection durability^8^.

The 2022 MPXV variants causing the outbreak form the lineage B.1 branch that belongs to MPXV clade 3^9^. Because of high sequence identity, immunity elicited by attenuated vaccinia vaccine can provide protection from monkeypox and smallpox. Both MPXV and vaccinia virus belong to Orthopoxvirus genus, which also includes variola virus, the pathogen of smallpox. Proteomic analysis of antibody response to MPXV infection and vaccinia vaccination revealed a number of immunodominant envelope proteins^10,11^. Among them, vaccinia A27L^12^, D8L^13^, H3L^14^, L1R^15^, A33R^16^ and B5R^17^ have been identified as targets of neutralizing antibodies. Polyvalent DNA vaccines combining vaccinia antigens A27L, D8L, L1R, A33R and B5R protected macaques and mice from lethal MPXV challenge^18,19^. The MPXV equivalent antigens of vaccinia A27L, D8L, L1R, A33R and B5R are A29L, E8L, M1R, A35R and B6R respectively. A number of important questions arise when exploring opportunities of developing a MPXV based vaccine. For example, how different are the MPXV and MVA antigens? Could an mRNA vaccine against MPXV elicit potent antibody response? Would an mRNA vaccine encoding these antigens be generally safe?

The mRNA vaccine technology has catalyzed the tremendous success of COVID mRNA vaccine, which has proven to be safe and effective across the globe in both initial clinical trials and real-world data. It is a versatile platform that can be adopted to develop mRNA vaccines against other infectious diseases, including monkeypox. Although MVA-based vaccine has less side effect than ACAM2000, its >=3 grade adverse event rate is 7.7% in recipients^6^. In contrast, the >=3 grade adverse event rate of COVID mRNA vaccine is similar to placebo group (1.5% in vaccine group vs. 1.3% in placebo group)^20^. The MVA vaccine contains around 200 proteins, many of which are not immunogenic and likely associated with undesired side effects. Removal of undesired components in MPXV vaccines would improve its safety profile, which is the case for MVA that lost large fragments of genome during attenuation^21^. Simplification of MPXV vaccine could facilitate rapid vaccine optimization. In addition, the manufacturing of mRNA vaccine can be achieved in vitro independent of complex cell culture as opposed to inactivated or attenuated vaccine; and is easily scalable by biochemical reactions. These features together make mRNA vaccine promising to enable the world to quickly fill the gap between vaccine supply and demand during a disease outbreak. The shortage of monkeypox vaccine and adaptability of mRNA vaccine prompt the idea of developing MPXV-targeting mRNA vaccine. However, whether the mRNA-based MPXV antigens can elicit significant immune response in vivo remains a critical question to be answered.

We designed a polyvalent mRNA vaccine candidate, MPXVac-097, against 2022 MPXV and initiated the testing of its antibody response as well as T cell receptor repertoire in vaccinated mice. Five MPXV antigens with prior evidence as neutralizing antibody targets, A29, E8L, M1R, A35R and B6R, were used in this vaccine design (**Fig. 1a**). They were linked in tandem by 2A peptides and codon optimized based on the protein sequence of 2022 MPXV case identified in USA in May, 2022^22^. Their sequence alignment with homologs of MVA showed over 90% sequence identity (**Supplementary Table 1**), and identified dozens of species-specific residues (**Fig. 1a and supplementary Fig. 1**). M1R antigen is largely conserved between MPXV and MVA, while the other four antigens contain 4-6% substitutions. Among these substitutions, K149E in A35R and T144A, S65T, L66I, R67H in E8L are located on reported neutralizing antibody interface^23,24^ and may lead to immune evasion of MVA neutralizing antibodies (**Fig. 1b**). The MPXV mRNA vaccine adopts the circulating MPXV sequence and can avoid this antigen mismatch. Interestingly, to prevent host cell interaction, the neutralizing antibody of E8L recognizes and blocks the ligand binding site which contains a large positive-charge patch to accommodate the negative charged chondroitin sulfate ligand of host cells (**Supplementary Fig. 2**).

**Fig. 1.**
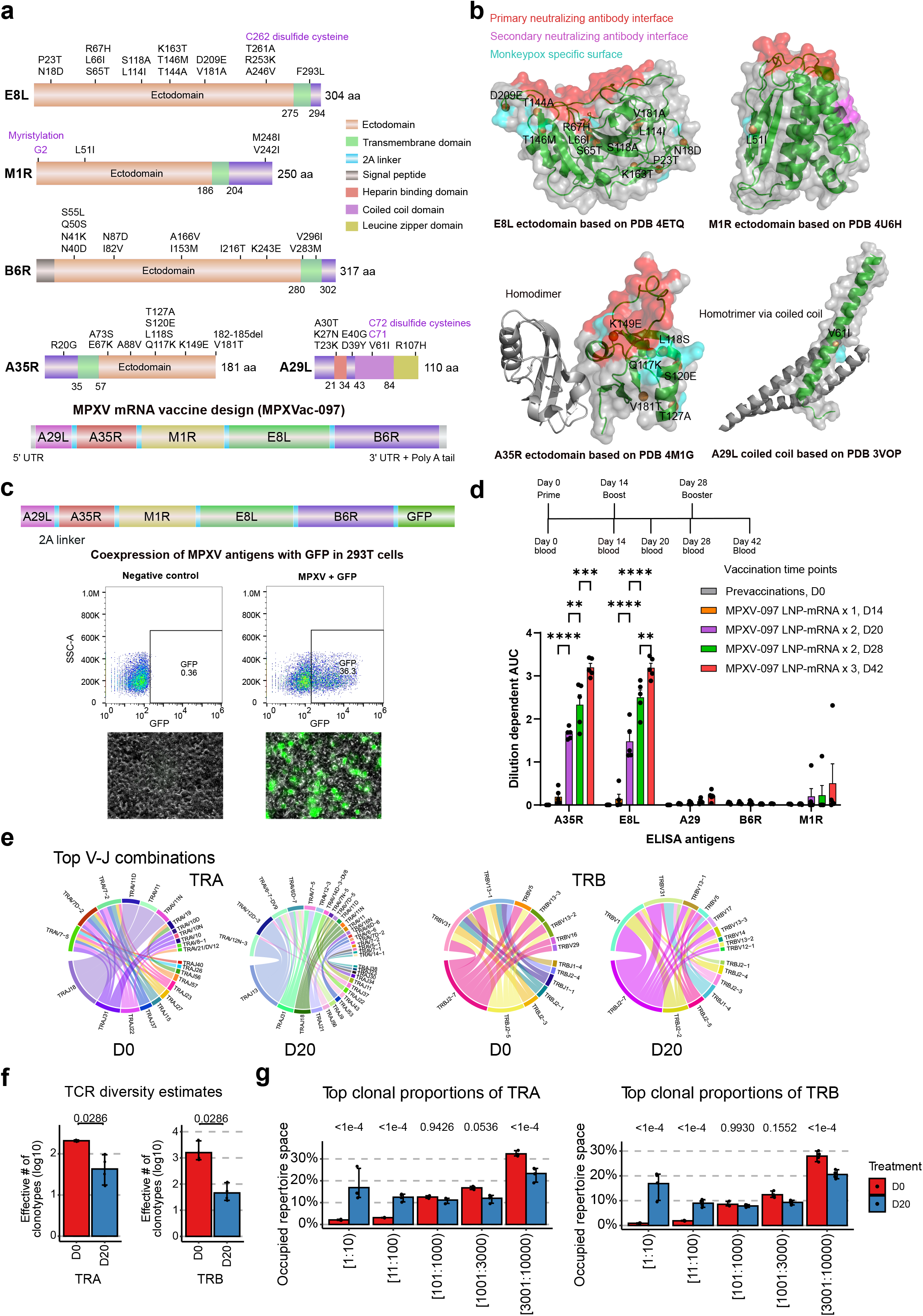
Monkeypox mRNA vaccine design and immunogenicity characterization in mice. **a**, Five neutralizing targets of monkeypox mRNA vaccine candidate are connected by 2A linkers and translated from the same mRNA transcript. Individual monkeypox antigen domain arrangement is shown along with antigen sequence difference between monkeypox and modified vaccinia ankara (MVA, attenuated monkeypox vaccine progenitor). The antigen design of a multivalent monkeypox LNP-mRNA vaccine candidate, MPXVac-097, is shown in the bottom panel. **b**, The antigen sequence differences between monkeypox and MVA are displayed on corresponding homology structural models. The reported neutralizing antibody epitopes on antigen surface are colored in red/magenta and monkeypox specific surface that differs from MVA is colored in cyan. **c**, Co-expression of five monkeypox antigens and GFP in 293T cells. The GFP positive cells with expression of five monkeypox antigens are identified by flow cytometry and fluorescence microscope. **d**, Three doses of MPXVac-097 LNP-mRNA significantly increased antibody titers against A35R and E8L (n = 5). Mice vaccination and blood collection schedule are illustrated on the time axis. Statistical comparisons were made between adjacent time points. **e**, Circos plots of the top 20 TRA and TRB V-J gene combinations for pre-vaccination (D0) and post-boost (D20) of two-dose MPXVac-097 vaccinated mice. replicate-pooled count data are presented in each plot. **f**, Bar plots of the estimated TCR diversity in TRA and TRB repertoires between pre-vaccination (D0) and MPXV post-boost (D20) mice. Estimates were calculated using the “true diversity” method, based on amino acid sequences. **g**, Bar plots of the relative distribution of TCR clones between different ranges of top clonotypes. The data are presented with the index range of the most abundant clonotypes on the x axis and the percent of total repertoire on the y axis. Fig. 1e-1g analyses were performed with n = 4 paired pre-vaccination and post-boost samples. Data on bar plots are shown as mean ± s.e.m. with individual data points in plots.

To test whether this multivalent vaccine construct encoding five monkeypox antigens in tandem could be successfully translated, a reporter construct otherwise identical to the vaccine was made in parallel by appending GFP to the end of tandem construct (MPXV-PentAg-GFP) (**Fig. 1c**). After transfection with this GFP reporter, around 36% of 293T cells expressed GFP which was quantified by flow cytometry (**Supplementary Fig. 3**) and observed under fluorescence microscope. The expression of GFP represents the successful translation of all residues from same mRNA transcript in a significant population of cells. We then advance the MPXVac-097 mRNA expression construct to lipid nanoparticle (LNP) formulation (**Methods**). The transcribed MPXVac-097 mRNA was successfully encapsulated in LNP, of which size distribution was determined by dynamic light scattering. The encapsulated MPXVac-097 LNP mRNA was monodispersed with an average radius of 49 nm and polydispersity index of 0.16 (**Supplementary Fig. 4**).

After confirming the successful translation of tandem construct and uniform size of the MPXVac-097 LNP-mRNA vaccine candidate, we characterized its initial immunogenicity in mice. Mice were immunized with three doses of 8 μg MPXVac-097 LNP mRNA two weeks apart (prime, boost and booster). The retro-orbital blood was collected on day 0, 14, 28 (just before prime, boost, booster) and day 20, 42 (6 days post boost and 14 days post booster). The isolated plasma from blood was used to quantify antibody titers against individual monkeypox antigen in enzyme-linked immunosorbent assay (ELISA). Two weeks post prime on day 14, a modest and insignificant increase of antibody titers against A35R and E8L was found in half of vaccinated mice (**Fig. 1d and supplementary Fig. 5**), suggesting a single dose (prime alone) is insufficient to elicit a strong antibody response. After the second and third doses, the antibody titer of A35R and E8L increased substantially, and became evident and significant in all mice on day 20, 28 and 42 (**Fig. 1d and supplementary Fig. 5**). Interestingly, the antibody titer against M1R, that had no change post prime, started to increase moderately post boost and booster in one mouse (**Fig. 1d and supplementary Fig. 5**). Because the M1R antigen was sandwiched between A35R and E8L on the same mRNA transcript, its expression level would be comparable to A35R and E8L. The delayed increase of M1R antibody titer in one mouse was unlikely due to differential expression of antigens and may stem from the difference of antigen immunogenicity or surface display. Minimal antibody response was detected for either A29L or B6R post boost, again highlighting the differences in immunogenicity between these antigens in the setting of mRNA vaccination.

The antigens used in ELISA are purified ectodomains or truncating variants. Next, we sought to ask if the full-length MPXV antigens can be presented on cell surface and recognized by the plasma antibodies elicited by MPXVac-097. The five monkeypox antigens were separately expressed in 293T cells from which surface antigens were detected by plasma antibody from mice on day 20 and anti-mouse IgG antibody with PE on flow cytometry (**Supplementary Fig. 6**). Expression of A35R and E8L led to an increase (12.6% and 1.2% respectively vs. 0.4% in negative control of SARS-CoV-2 spike) of 293T cells that were bound to the MPXV plasma antibodies (**Supplementary Fig. 6**). This increase of MPXV antibody-bound cells was consistent with findings in ELISA, and validated the observation of antibody response to surface presentation of A35R and E8L cellular antigens.

The ELISA and cell-surface antigen assays provided evidence of the initial B cell response to five monkeypox antigens in mice immunized with MPXVac-097. A natural question that follows is whether T cell response was induced in these animals. We went on to characterize the T cell response by profiling the repertoire of T cell receptors (TCR) in vaccinated mice via bulk TCR-sequencing (TCR-seq). TCR-seq of peripheral blood mononuclear cells (PBMCs) revealed VJ combination usage and clonality maps of TCRs from mice before and after two MPXVac-097 LNP mRNA injections on day 0 and day 20, respectively (**Fig. 1e-1g; Supplementary Fig. 7-9**). MPXVac-097 LNP mRNA vaccination resulted in more frequent use of TRAJ13, TRAV12N-3, TRAV12D-3 in alpha chain and TRBJ1-4, TRBJ2-2, TRBV1 in beta chain (**Fig. 1e**). After immunization, the most frequent VJ combinations was switched to TRAJ13-TRAV12N-3/TRAV12D-3 and TRBJ2-7-TRABV1. The total number of clones and unique clones were not significantly different between samples on day 0 and 20 (**Supplementary Fig. 8b**). The CDR3 length distributions are also similar (**Supplementary Fig. 9a**). In contrast, while the TCR repertoire compositions are similar between independent mice pre-vaccination, we observed distinct TCR repertoire compositions between day 0 and day 20 groups, suggesting that repertoire compositions changed substantially upon two-dose MPXVac-097 vaccination (**Supplementary Fig. 9b-c**). Moreover, the clonal diversity of samples post boost was significantly reduced in two-dose MPXVac-097 vaccinated as compared to vaccine-naïve animals (**Fig. 1f**). There were also significant reduction of low-frequency clones and significant increase in expended or hyperexpanded clones (**Fig. 1g and Supplementary Fig. 9d**), which together are indicative of clonal expansion induced by MPXVac-097 vaccination.

In summary, our study designed a multivalent monkeypox LNP-mRNA vaccine candidate, MPXVac-097, which can elicit significant antibody response against a subset of MPXV antigens (A35R and E8L) as well as T cell response in mice. Taken together, these data demonstrated initial feasibility of MPXV mRNA vaccine design and provided initial evidence of functionality for its future optimization.

## Supplementary information

### Supplementary Figure Legends

**Supplementary Figure 1.**
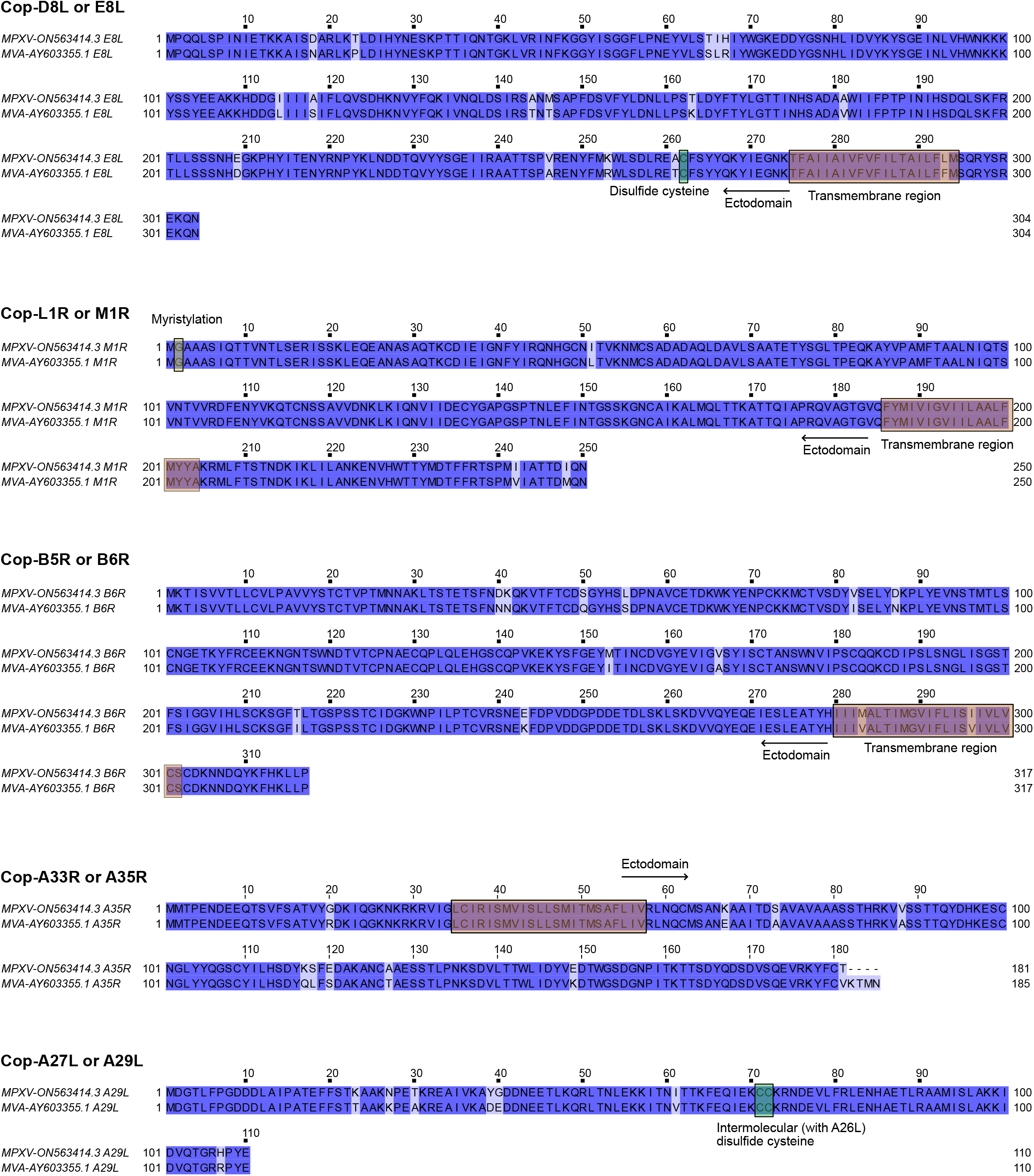
Sequence alignment of five antigen homologs from monkeypox (GenBank ON563414.3) and MVA (GenBank AY603355.1). Transmembrane domains and ectodomains of each antigen are indicated on the sequences.

**Supplementary Figure 2.**
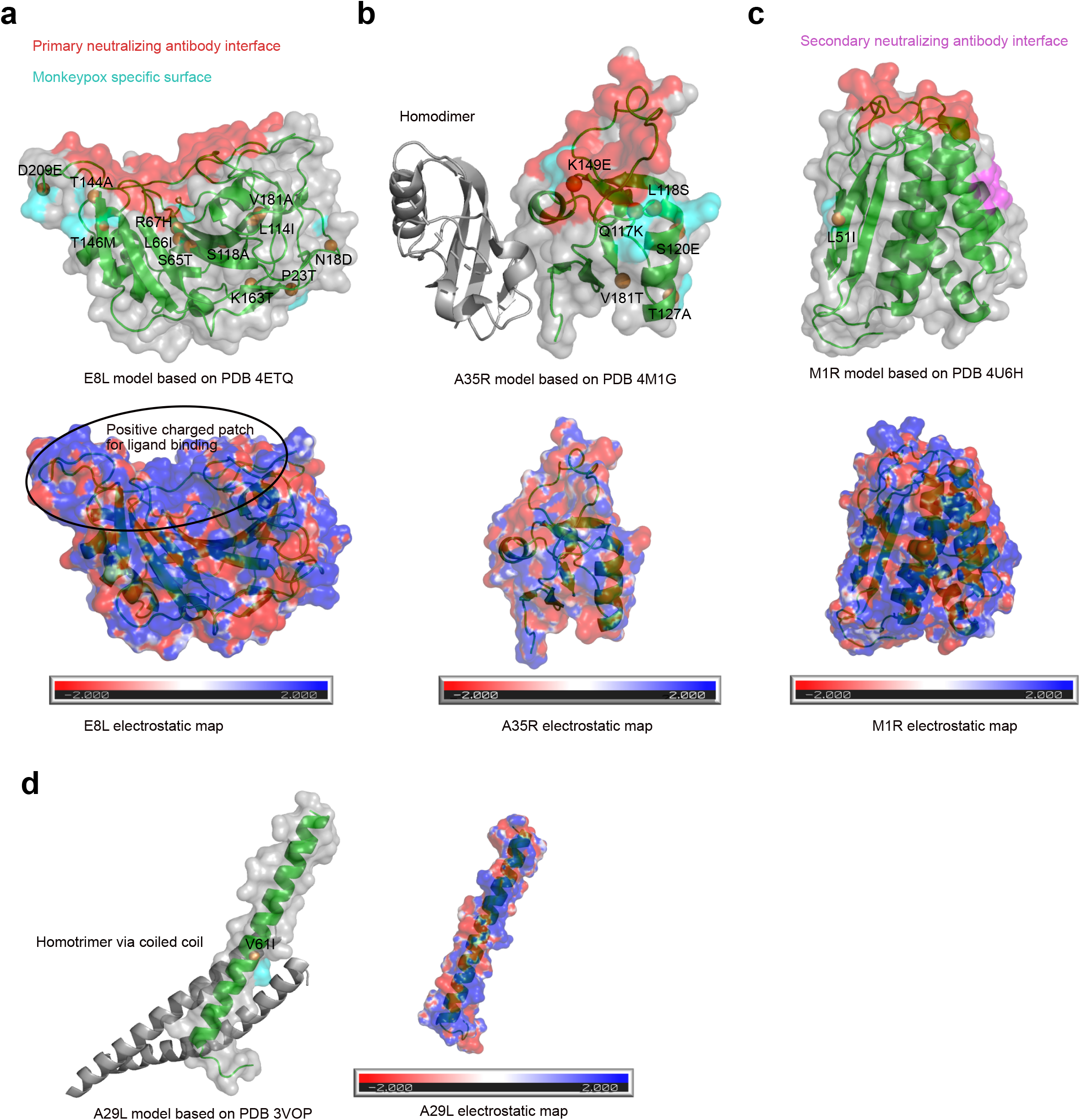
Calculated electrostatic map on homolog structures of five monkeypox antigens (a-d). The surface electrostatic map was generated using CHARMM-GUI server (<u>CHARMM-GUI</u>) and SWISS homology models (<u>SWISS-MODEL (expasy.org)</u>).

**Supplementary Figure 3.**
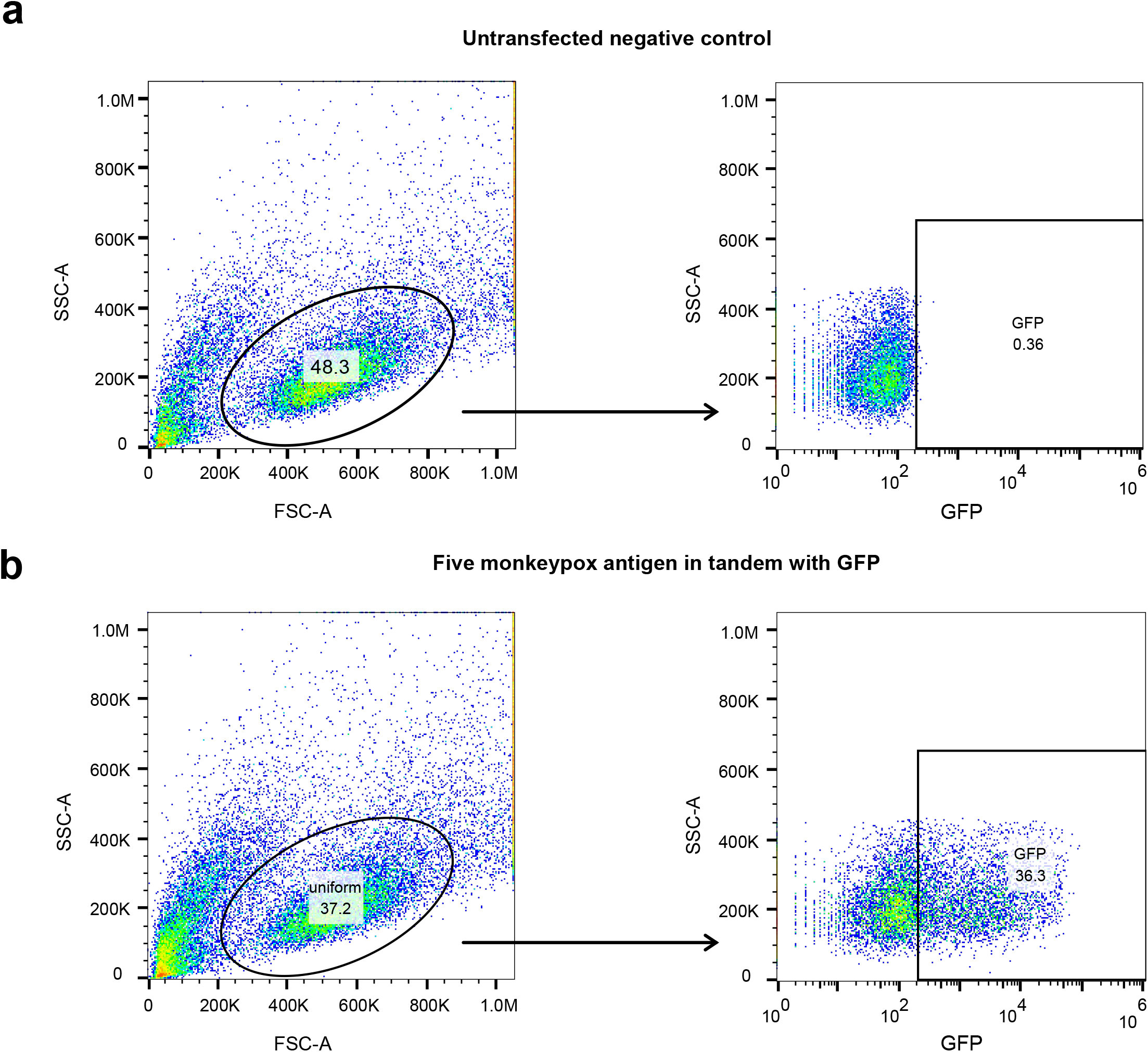
Untransfected 293T cells as negative controls in the flow cytometry and fluorescence microscope experiments. **a**, gating strategy used in flow cytometry.

**Supplementary Figure 4.**
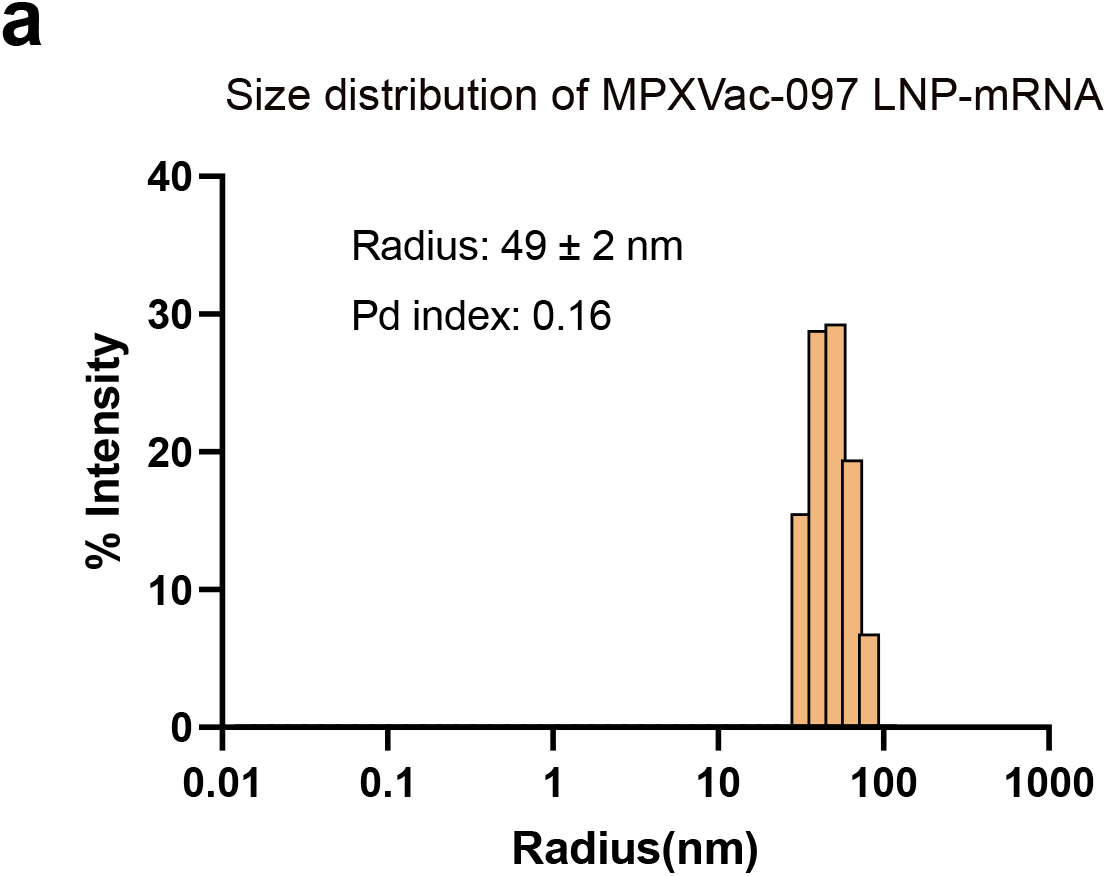
Size distribution of MPXVac-097 LNP-mRNA as determined by dynamic light scattering.

**Supplementary Figure 5.**
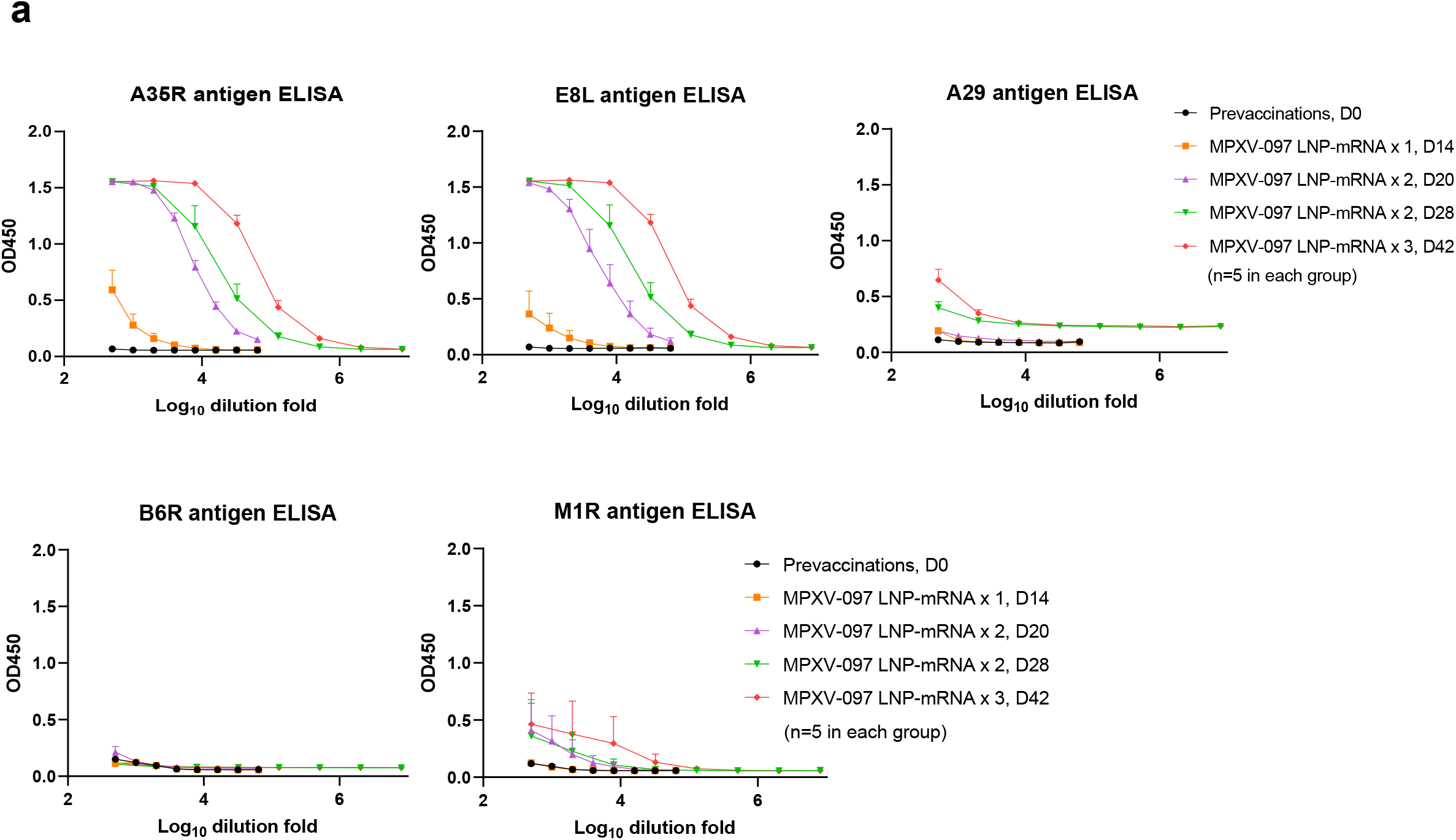
ELISA titration curves showing binding response (OD450 on y axis) over serial dilution points of plasma samples (log_10_ transformed dilution factor on x axis) collected from mice vaccinated with zero, one, two or three doses on MPXV LNP-mRNA on day 0, 14, 20, 28 and 42.

**Supplementary Figure 6.**
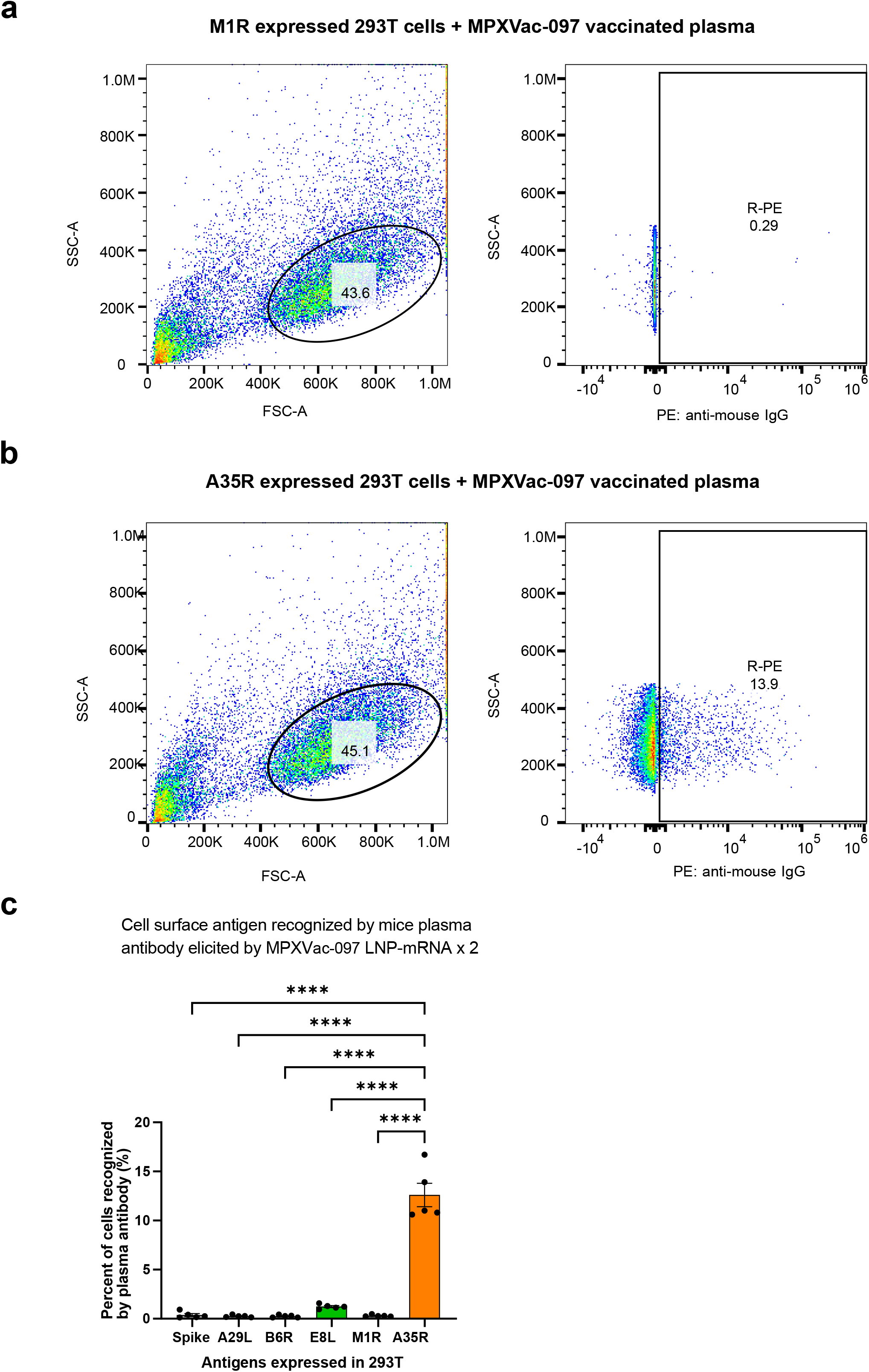
Flow cytometry to identify mice antibody bound 293T cells expressing monkeypox antigens. Plasma samples were collected on day 20 from mice immunized with two doses of MPXV LNP-mRNAs and the primary mice antibody bound to the cells was recognized by anti-mouse IgG second antibody with PE fluorophore. **a-b**, gating strategy of flow cytometry for cells expressing M1R (a) or A35R (b) antigens. **c**, Monkeypox antigens expressed on 293T cell surface were recognized by plasma antibody produced from mice on day 20. SARS-CoV-2 spike expressing cells served as a negative control. Data on bar plots are shown as mean ± s.e.m. with individual data points in plots.

**Supplementary Figure 7.**
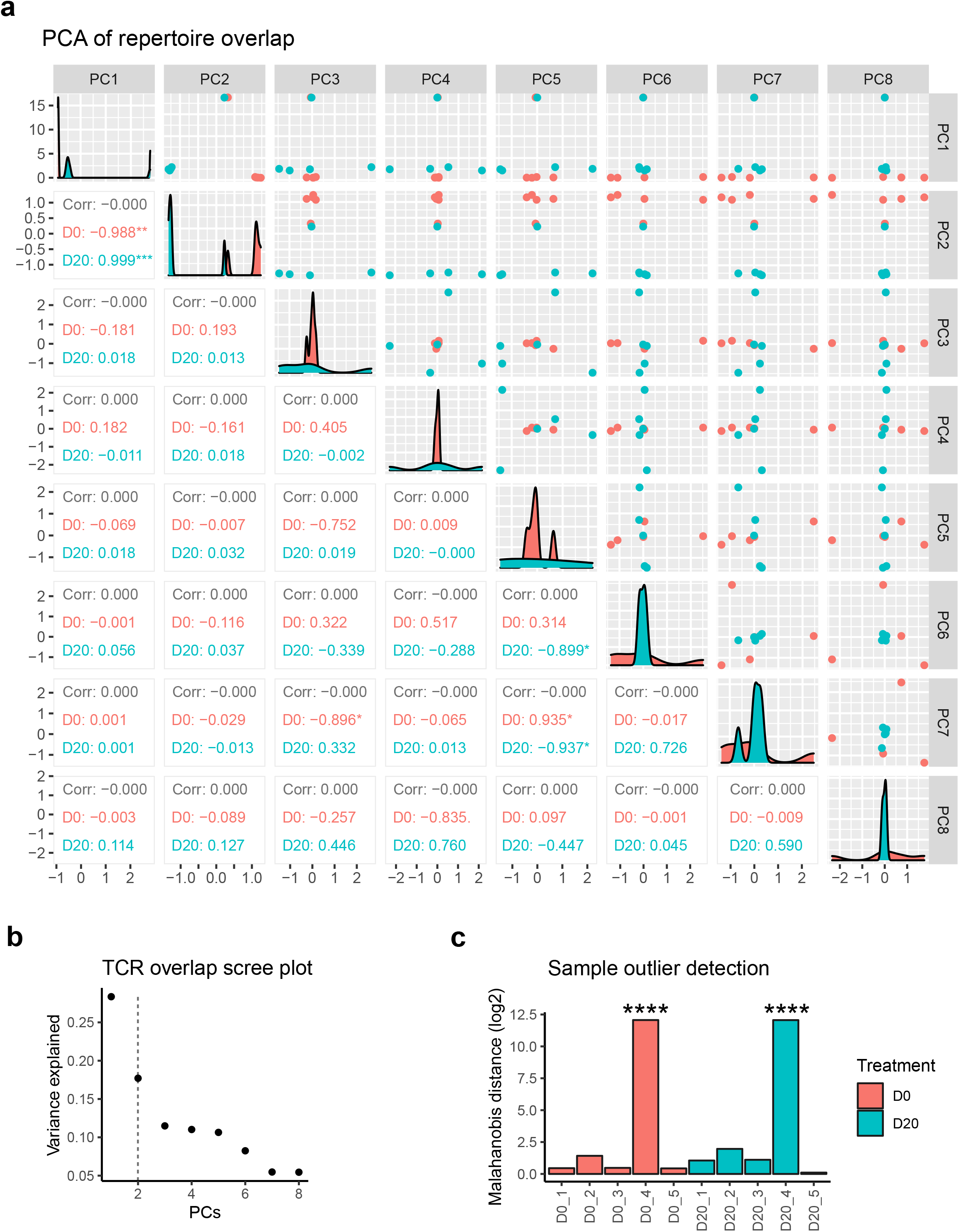
Quality control and sample filtering of TCR sequencing in pre-vaccinated and MPXV post-boosted mice. **a**, Plots comparing the first eight components from the principal components analysis (PCA) of TCR repertoire overlap between samples. TCR repertoire overlap (amino acid sequence with variable region gene) was assessed by Morisita index score for TRA and TRB separately, and PCA was performed with pooled index scores. Plots in the upper-right triangle compare PCs, while the panels of the bottom-left triangle show Pearson correlation rho for the corresponding upper-right panel. Correlation rho are provided for total, pre-vaccination (D0), and post-boost (D20) in black, red, and blue, respectively. **b**, Scree plot of the variance explained by each PC from the PCA of TCR repertoire overlap between samples. First two PCs were chosen for further analysis based on the elbow plot method. **c**, Bar plot of the sample differences, using Malahanobis distance calculated from the first two PCs. Statistical analyses were performed by chi-squared test for each sample, and results were FDR-corrected for multiple testing. Statistical significance labels: * p < 0.05; ** p < 0.01; *** p < 0.001; **** p < 0.0001. Non-significant comparisons are not shown.

**Supplementary Figure 8.**
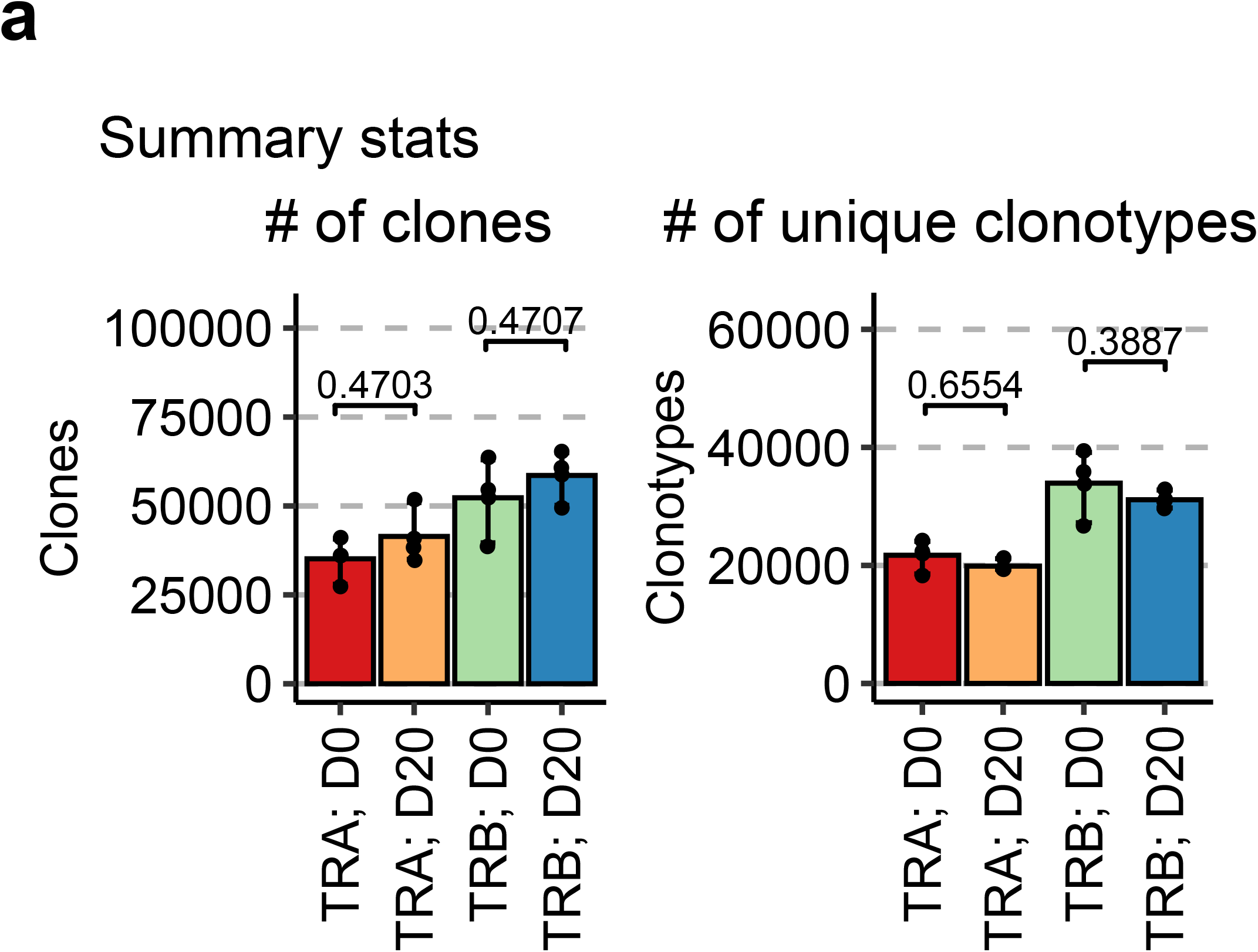
TCR gene repertoire analyses in pre-vaccinated and MPXV post-boosted mice. **a**, Bar plots for the number of clones and unique clonotypes detected between pre-vaccination (D0) and MPXV post-boost (D20) mice. All analyses were performed with n = 4 paired pre-vaccination and post-boost samples. Data on bar plots are shown as mean ± s.e.m. with individual data points in plots.

**Supplementary Figure 9.**
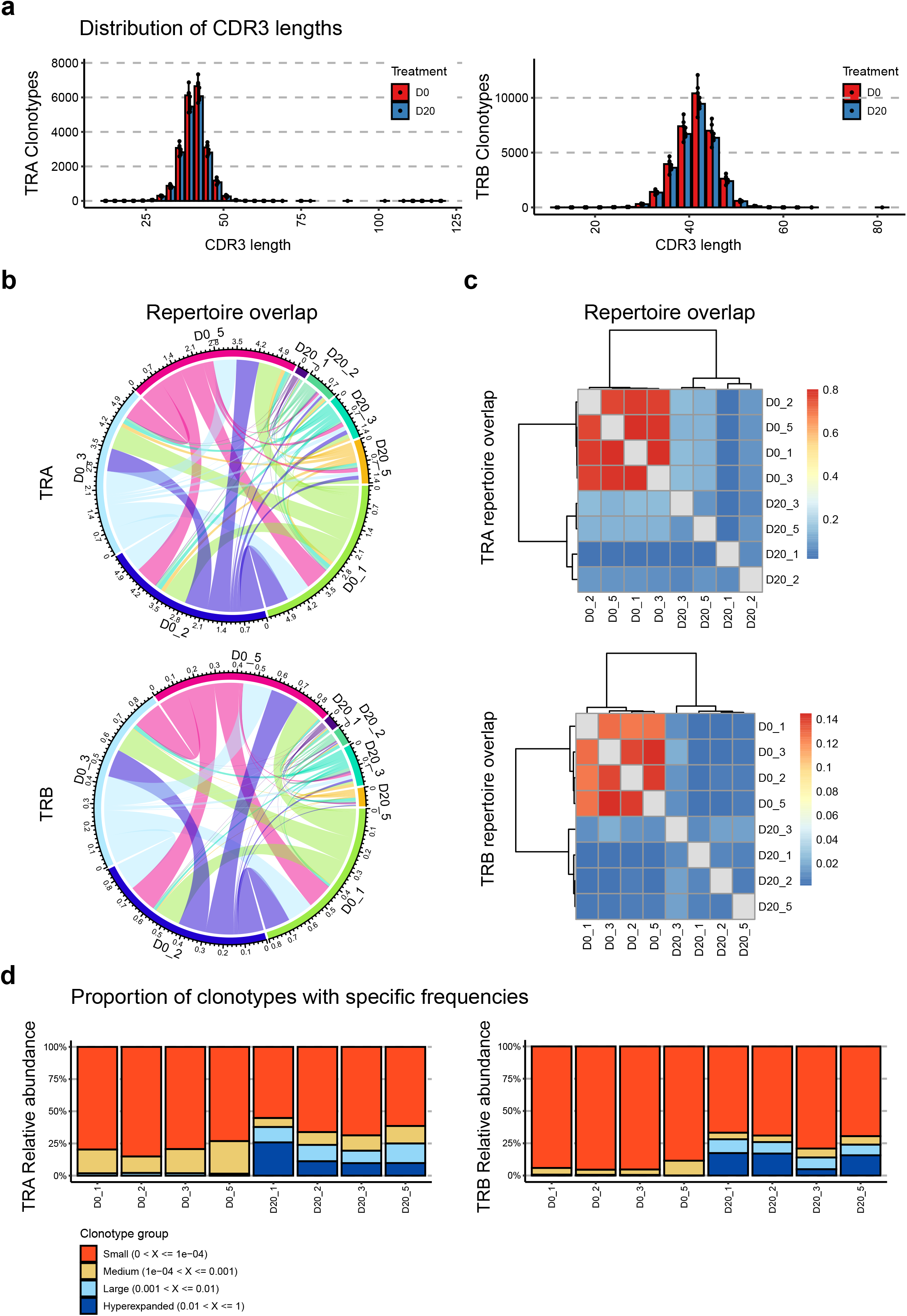
Summary statistics of TCR sequencing in pre-vaccinated and MPXV post-boosted mice. **a**, Histograms of the CDR3 amino acid lengths for TRA and TRB clonotypes between pre-vaccination (D0) and post-boost (D20) two-dose MPXVac-097 vaccinated mice. **b-c**, Circos, and bar plots (**b**, and **c**, respectively) of the TRA and TRB repertoire overlap between pre-vaccination (D0) and MPXV post-boost (D20) mice. Repertoire comparisons were performed using the Morisita method. **d**, Stacked bar plots of the relative abundances of clones between different ranges of clonotype frequencies. The data are presented with the index range of the most abundant clonotypes on the x axis and the percent of total repertoire on the y axis.

## Methods

### Institutional approval

The work described in this study is performed under relevant approved protocols (IACUC, rDNA/EHS) in place.

### Plasmid cloning

The monkeypox antigen amino acid sequences used in the mRNA vaccine design were based on the genome of confirmed monkeypox case identified in Massachusetts in May, 2022 (Genbank ON563414.3)^22,25^. The five antigen cDNAs were codon optimized, synthesized and inserted to mRNA vector with 5’, 3’ UTRs and poly A tail as flanking sequences. The MPXVac-097 mRNA vaccine vector was cloned through connecting five antigen cDNAs with 2A linkers as shown in **Fig. 1a**. A reporter vector with GFP appended to the 3’ end of the mRNA vaccine vector, MPXV-PentAg-GFP, was designed to validate the successful translation of mRNA in mammalian cells (**Fig. 1c**).

### SWISS homology modeling

Homologs of four out of five monkeypox antigens were found in PDB database and were used to build the homology structures on SWISS homology-modeling server (<u>SWISS-MODEL (expasy.org)</u>). The homology model was visualized in Pymol and was used to display neutralizing antibody epitopes and sites that are different between monkeypox and MVA.

### Cell Culture

HEK293T cells (ATCC, CRL-3216) were cultured in Dulbecco’s modified Eagle’s medium (DMEM, Thermo fisher) with 10% hea-inactivated fetal bovine serum (FBS, Hyclone). Cells were split every other day at a split ratio of 1:4 or when confluency reaches over 95%.

### Monkeypox antigen expression in 293T cells

The Lipo3000 transfection system (Thermo Fisher, Cat. No. L3000015) was applied to transfect 293T cells with a reporter vector that contains five monkeypox antigens in tandem and a 3’-end GFP. The 293T cells were seeded on the 6-well plates one day before transfection and transfection was performed according to manufacture’s protocol. Two days after transfection, cells were imaged under fluorescence microscope and were subsequently subject to flow cytometry for quantification of cells co-expressing five monkeypox antigens and GPF.

### Cell surface monkeypox antigen staining using mice plasma antibody

The same lipo3000 transfection approach was also applied to express individual monkeypox antigen in 293T cells. The 293T cells expressing monkeypox antigen were trypsinized, washed with 2% FBS in PBS once and resuspended in flow cytometry staining buffer (Thermo Fisher, Cat. No. 00-4222-26). Resuspended cells were incubated on ice for 20 min with 1:100 diluted plasma from mice vaccinated with two doses of MPXV LNP-mRNA. After incubation, cells were washed once with flow cytometry staining buffer and incubated on ice for 20 min with 1:100 diluted anti-mouse IgG antibody conjugated to PE fluorophore (Thermo Fisher, Cat. No. P-852). After the final wash with flow staining buffer, PE-positive cells were detected by flow cytometry using Attune focusing cytometer (Attune NxT Software v3.1). FlowJo v.10.7.1 was used for flow cytometry data analysis.

### In vitro mRNA transcription and lipid nanoparticle preparation

To produce mRNA, the MPXVac-097 vaccine candidate construct encoding five monkeypox antigens in tandem was in vitro transcribed from linear DNA template using HiScribe T7 ARCA mRNA kit (NEB, Cat. No. E2060S) with 50% replacement of uridine by N1-methyl-pseudouridine (Tirlink, Cat. No. N-1081-5). The transcribed mRNA was purified using Monarch RNA Cleanup Kits (NEB, Cat. No. T2040L) and kept in - 20 until further use.

The prepared mRNA was diluted in 25mM sodium acetate buffer at pH 5.2 and mixed with lipid mixture in ethanol using NanoAssemblr^@^ Ignite™ instrument (Precision Nanosystems). The lipid mixture is composed of ALC-0315, ALC-0159, DSPC and cholesterol at a mixing ratio as previously described^26^. The MPXV LNP-mRNA buffer was exchanged to phosphate buffered saline (PBS) using 100kDa Amicon filter (Macrosep Centrifugal Devices 100 K, 89131-992). The mRNA encapsulation rate and mRNA concentration were determined by Quant-iT™ RiboGreen™ RNA Assay (Thermo Fisher). The size distribution of LNP-mRNA sample was characterized by dynamic light scattering (DynaPro NanoStar, Wyatt, WDPN-06). Sucrose was added to the final LNP-mRNA sample before storing it in -80 freezer.

### Mouse immunization

8-10 week-old female C57BL/6Ncr mice purchased from Charles River were vaccinated with two doses of 8 μg MPXVac-097 LNP-mRNA vaccine candidate on day 0 (prime), day 14 (boost) and day 28 (booster). The mice were maintained at an ambient room temperature, 40-60% humidity and a 14h:10h day/night cycle. Retro-orbital blood was drawn using heparinized micro capillary tubes (Fisher, Cat. No. 22362566) and collected in lithium heparin microtainers (Fisher, Cat. No. 13-680-62) just before prime, boost and booster on day 0, 14 and 28. Additional blood samples were collected in the same way on day 20 (6 days post boost) and 42 (14 days post booster).

### Isolation of plasma and peripheral blood mononuclear cells (PBMCs) from blood

The collected blood was 1:1 diluted with 2% FBS and was added to 7ml of Lymphoprep™ density gradient medium in SepMate-15 tubes (StemCell Technologies). The red blood cells, PBMCs and plasma were isolated from blood by centrifugation at 1200 x g for 20 min. After centrifugation, approximately 200 to 300ul plasma was collected from the surface layer and the PBMCs in the remaining solution at the top were poured to a new tube. The separated PBMCs were washed with 2% FBS and sample’s mRNA was extracted using RNeasy Plus Mini Kit (Qiagen).

### ELISA

Commercial monkeypox antigens used in ELISA include E8L (AcroBiosystems, E8L-M52H3), M1R (Sino Biological, 40904-V07H), B6R (Sino Biological, 40902-V08H), A35R (Sino Biological, 40886-V08H), and A29L (Sino Biological, 40891-V08E). Recombinant antigens at 3 μg/ml in PBS were coated on the 384-well microplate (Fisher, Cat. No. 07-000-877) in cold room overnight. On next day, the plate was washed three time with PBST (0.05% Tween-20) on 50TS microplate washer (Fisher Scientific, NC0611021), and was blocked with 0.5% BSA in PBST at room temperature for one hour. Plasma was twofold serially diluted with PBS at a starting dilution of 1:500. The plasma diluent was added to the plate and incubated at room temperature for one hour. After washing with PBST five time, the plate was incubated with anti-mouse IgG (H+L) secondary antibody (Fisher, A16072) at room temperature for one hour. The secondary antibody with horse radish peroxidase was 1:2500 diluted in 0.5% BSA blocking buffer before adding to the microplate. The plate was washed five times with PBST and developed with tetramethylbenzidine substrate (Biolegend, 421101). After 20 min at room temperature, the reaction was stopped with 1M phosphoric acid and OD450 was measured by multimode microplate reader (PerkinElmer EnVision 2105, Envision Manager v1.13.3009.1401). The dilution-dependent area under curves (AUC) was calculated from AUC subtracted by the background AUC, which is the product of log dilution difference and end dilution OD450 (details in data summary excel file).

### Bulk TCR sequencing of PBMCs

Bulk TCR library was prepared using 200 ng mRNA extracted from PBMCs collected on day 0 (pre-vaccination) and day 20 (post boost) mice as described above. The SMARTer Mouse TCR a/b Profiling Kit (Takara, Cat. No. 634403) was used to amplify both TCRα and TCRβ sequences. The purified amplicon concentration and purity was determined using D5000 high sensitivity tapes on TapeStation (Agilent). Equal amount of TCR amplicons from each sample was pooled and diluted with water to get a 2nM library. The pooled library was denatured, mixed with 5% PhiX and sequenced on MiSeq (Illumina) using MiSeq V3 2 × 300 cycle kit (Illumina).

### VDJ sequencing data analysis

The bulk VDJ sequencing data was pre-processed on MiSeq local run manager to trim adaptors and separate reads based on sample index. The reads with an average quality score less 30 were removed from downstream analysis. Clonotypes were called using MiXCR v2.1.5 with the recommended settings for 5’ RACE (RNA alignment to V gene transcripts with P region). The output was further processed with Immunarch v0.6.6 R package for statistical analyses using standard analysis pipelines. The initial samples were n = 5 independent mouse samples, assessed pre-vaccination and post-boost (labeled D0 and D20, respectively) in paired analyses. Samples were assessed for outliers by a multi-step process, (1) comparing the repertoire overlap (amino acid sequence and V gene) between samples using the Morisita method, separately for TRA AND TRB, (2) performing principal components analysis on combined TRA and TRB overlap information (Morisita indices), (3) determining the optimal PCs for subsequent analysis (details below), (4) calculating Mahalanobis sample distances using the optimal number of PCs, and (5) calculating p values using a chi-squared distribution (2 degrees of freedom for 2 PCs) and correcting for multiple testing using the FDR method (**Supplementary Figure 7a-c)**. The optimal number of PCs was chosen as two using the elbow plot method, and the choice was supported by PC scatter plots that demonstrated that PC1 best explained outlier samples, PC2 best explained the treatment differences, while Pearson correlation analyses of PC1 and PC2 demonstrated the best treatment specific correlation. The filtering parameter excluded two TCR libraries, one from each group (D0 or D20), and the final TCR analysis consists of four samples from each group (n = 4 each).

### Statistics and Reproducibility

Standard statistical methods were applied to experimental data. The statistical methods are described in here, figure legends and/or supplementary Excel tables. Data on dot-bar plots are shown as mean ± s.e.m. with individual data points in plots. Two-way ANOVA with Tukey’s multiple comparisons test was used to assess statistical significance for grouped datasets in **Fig. 1. Fig. 1’s** statistics were calculated using a 2-way ANOVA test with Sidak’s multiple test comparison for panel g, and a Kolmogorov-Smirnov test for **f**. Statistical significance labels: * p < 0.05; ** p < 0.01; *** p < 0.001; **** p < 0.0001. Non-significant comparisons are not shown, unless otherwise noted as n.s., not significant. Sample number is designated as n from biologically independent samples. Prism (version 9.3.2, GraphPad Software Inc.) and RStudio (version 1.3.959, RStudio software) were used for these analyses. Additional information can be found in the supplementary excel tables.

### Schematic illustrations

Schematic illustrations were created with Affinity Designer or BioRender.

### Replication, randomization, blinding and reagent validations

In animal experiments, mice were randomly assigned to each cage or vaccination group. Experiments were not blinded.

Commercial antibodies were validated by the vendors, and re-validated in house as appropriate. Cell lines were authenticated by original vendors, and re-validated in lab as appropriate. All cell lines tested negative for mycoplasma.

## Data availability

All data generated or analyzed during this study are included in this article and its supplementary information files. Specifically, source data and statistics for regular experiments are provided in an excel file of Source data and statistics. NGS data are being deposited to SRA/GEO with pending accession codes. Other materials and data are available either through public repositories, or via reasonable requests to the corresponding authors to the academic community.

## Code availability

Custom codes are available either through public repositories or via reasonable requests to the corresponding authors to the academic community.

## Reporting summaries

### Statistics

For all statistical analyses, we confirmed that the items mentioned in NPG reporting summary are present in the figure legend, table legend, main text, or Methods section.

### Standard statistical analysis

All statistical methods are described in figure legends, methods and/or supplementary Excel tables. Source data and statistics were provided in a supplemental excel table.

### Software and code

#### Data collection

Default softwares in the data collection instruments including Attune focusing cytometer (Attune NxT Software v3.1) and PerkinElmer EnVision 2105 microplate reader (Envision Manager v1.13.3009.1401) were used to collect data.

#### Data analysis

Data analysis were performed using Prism (version 9.3.1, GraphPad Software Inc.) for small scale data. NGS data were analyzed with custom codes.

#### Code availability

Custom codes are available either through public repositories, or via reasonable requests to the corresponding author to the academic community.

#### Life sciences study design

#### Sample size determination

For most cases, each group has five biologically independent samples unless otherwise noted. Details on sample size for each experiment were indicated in methods and figure legends. Sample size was determined according to the lab’s prior work or similar studies in the field.

#### Data exclusions

Five independent mice were initially used for TCR-seq library prep. In two TCR libraries, one paired sample No. 4 from two groups (D0 or D20) did not pass QC and thus were excluded from subsequent analysis. This is noted in methods section and figure legend.

No other data were excluded.

#### Replication

Biological replicates of mouse samples were done in each experiment.

#### Randomization

Mice were randomly allocated into experimental or control groups.

#### Blinding

The experiments were not blinded.

## Reporting for specific materials, systems and methods Antibodies used

1. HRP-conjugated anti-mouse IgG (H+L) antibody (Fisher, Cat. No. A16072, 1:2500 dilution)
2. PE-conjugated anti-mouse IgG (H+L) antibody (Fisher, Cat. No. P-852, 1:100 dilution)

### Antibody validation

Commercial antibodies were validated by the vendors, and re-validated in house as appropriate through antigen-specific experiments. Commercial antibody info and validation info where applicable:

1. HRP-conjugated anti-mouse IgG (H+L) antibody (Fisher, Cat. No. A16072, 1:2500 dilution): <u>Goat anti-Mouse IgG (H+L) Cross-Adsorbed, HRP (A16072) (thermofisher.com)</u>
2. PE-conjugated anti-mouse IgG (H+L) antibody (Fisher, Cat. No. P-852, 1:100 dilution): <u>Goat anti-Mouse IgG (H+L) Cross-Adsorbed, PE (P-852) (thermofisher.com)</u>

### Eukaryotic cell lines

Commercial cell line HEK293T was originally acquired from ATCC (CRL-3216).

### Authentication

Cell lines were authenticated by original vendors, and re-validated in lab as appropriate by morphology.

### Mycoplasma contamination

All the cell lines used here tested negative for mycoplasma contamination.

### Commonly misidentified lines (See ICLAC register)

No misidentified cell lines were used in the study.

### Animals and other organisms Laboratory animals

C57BL/6Ncr (B6) mice, female 8-10 week mice purchased from Charles River.

### Wild animals

N/A

### Field-collected samples

N/A

### Flow Cytometry Plots

Confirm the checkboxes of requirements.

### Methodology

#### Sample preparation

Various sample prep details are provided in the Methods section

#### Instrument

Flow cytometry data were collected on Attune focusing cytometer (Attune NxT Software v3.1)

#### Software

FlowJo v.10.7.1 was used for flow cytometry data analysis.

#### Cell population abundance

N/A

#### Gating strategy

Cells were gated by FSC/SSC plot. To distinguish between positive and negative boundaries of the stained cells, negative control samples were analyzed and utilized as background.

#### Gating example figure

We confirm that a figure exemplifying the gating strategy is provided in the Supplementary Information.

